# Warming affects routine swimming activity and novel odour molecular response in larval zebrafish

**DOI:** 10.1101/2023.06.16.545285

**Authors:** Jade M. Sourisse, Lucrezia C. Bonzi, Julie Semmelhack, Celia Schunter

## Abstract

Temperature is a primary factor affecting the survival, development, and physiology of aquatic ectothermic animals and global warming of water bodies may therefore impact several biological levels of aquatic life. Understanding the effects of near-future predicted temperature changes on the behaviour and the underlying molecular mechanisms of aquatic animals is of particular importance, since behaviour mediates key interactions and, in turn, population dynamics. In this study, we investigate the effects of elevated developmental temperature on locomotor behaviour and olfactory learning in the zebrafish, *Danio rerio*. We exposed zebrafish from cleavage embryonic stage to either current day control (28°C) or predicted future elevated temperature (30°C) for seven days. Overall, warming reduced the total routine swimming distance and caused the upregulation of a small number of genes involved in metabolism and neuron development, suggesting accelerated development at elevated temperature. When fish were exposed to two different olfactory cues, namely catfish cue, a non-alarming but novel odour, and injured conspecifics alarm cue expected to cause a fear reaction, warming differently affected larvae response to the two cues. In particular, a large transcriptional reprogramming was observed at elevated temperature in response to novel odour exposure, with upregulation of cell signalling, neuron development and neuron functioning genes. As this response was coupled with downregulation of genes involved in protein translation and ATP metabolism, it indicates that novel odour recognition in future-predicted thermal conditions will require energetic trade-offs between expensive baseline processes and responsive functions. To also evaluate their learning abilities at both temperatures, 7 days post fertilization (dpf) zebrafish were conditioned with a mixture of injured conspecifics alarm cue and non-alarming catfish cue. Regardless of temperature, no behavioural (freezing) nor gene expression changes were detected, reinforcing our previous findings that warming mainly affects zebrafish molecular response to novel odours. Overall, our results show that future thermal conditions will likely impact developing stages, causing energy trade-offs following olfactory detection of novel substances in the environment.

## Introduction

Olfaction is one of the main ways fish gather information about their environment. A wide variety of behaviours is influenced by the detection of olfactory stimuli, such as feeding, migration, reproduction and predator escape response (Hara, 1975). One famous example is the accidental discovery of fear reaction to alarm cues in fish: when a minnow (*Phoxinus phoxinus*) is injured, an alarm cue is released and nearby conspecifics drastically change their behaviour by swimming away and moving in tighter schools (von Frisch, 1938). Such olfactory-mediated behaviours are common in fish (Pfeiffer, 1977) and positively affect survival not only in adults but also in immature life stages, and many fish larvae can assess risks by sensing cues from predators or other prey (Wisenden, 2000). While some of the olfactory processes important for survival are innate, others involve associative learning: if during a conditioning phase two odours occur together, later in life the second odour will be associated with the same events and situations as the first odour (Suboski, 1990). This is the case with predator recognition, which has been described in the European minnow. After sensing the odour of a natural predator mixed with alarm cues released by the skin of conspecifics, minnows would start hiding and schooling upon smelling the pike’s odour alone (Magurran, 1989). Such an association of odours causing antipredator behaviours to otherwise neutral stimuli occurs in a diverse range of prey fishes (Brown, 2003), notably zebrafish (Jesuthasan & Mathuru, 2008).

Environmental factors have the potential to influence olfactory responses (Tigert & Porteus, 2023), with temperature arguably one of the main environmental variables influencing the performance of fish (Brett, 1971; Johnston & Dunn, 1987) including effects on olfactory triggered feeding (Stoner et al., 2006). Global warming, which is happening at an unprecedented rate in aquatic environments (Allen et al., 2018; Pachauri et al., 2014), may therefore alter crucial processes in fish, including olfactory and behavioural responses. Olfaction in fish is mediated by olfactory sensory neurons that converge on the olfactory bulb, where there is an exchange of information to second-order neurons, that are likely involved in the learning process (Laberge & Hara, 2001). Physiological changes in the olfactory bulb may occur due to temperature (Døving & Belghaug, 1977; Flerova & Gdovskii, 1975) and warming can impair fish associative learning due to alterations in memory formation (Toni et al., 2019; Závorka et al., 2020). Despite previous evidence of temperature altering gene expression involved in olfaction (Magnuson et al., 2023), it is unclear to what extent or how near-future thermal conditions will impact the sensory system and overall cue processing in fishes (Beltrán et al., 2021), particularly at the molecular level. For this reason, understanding how warming modulates the behaviour and learning experiences that fish need for survival is of crucial importance, especially in light of rapid climate change.

In our study, we investigate the effects of developmental temperature on the locomotory behavioural and transcriptomic responses of zebrafish larvae to two different olfactory cues: an alarm substance (Conspecifics Alarm Cue) and a non-alarming odour (catfish cue). We hypothesize that future predicted thermal conditions will impact the molecular state in sensory processing during the behavioural response to different cues. By performing classical conditioning, we moreover assess whether temperature influences the olfactory learning experience, both at behavioural and molecular levels. Overall, by characterizing temperature-specific transcriptomic and behavioural changes in response to olfactory cues, we aim at determining how future predicted temperature will affect the environmental perception of zebrafish and their learning experience through olfaction.

## Methods

### Animals, housing and temperature exposure

Wild-type breeders (nine females, ten males; AB strain) were obtained from the Hong Kong University of Science and Technology zebrafish husbandry facilities. Wild-type zebrafish were kept at 28°C and housed in two recirculating systems (80 × 37 × 32 cm) with six males/four females in one and four males/six females in the other, with a 14/10h light-dark cycle. The thermal conditions were measured, adjusted and recorded every 60 seconds with heaters (Schego) and a STC-1000 Thermostat (Elitech). Fish were fed twice a day TetraMin pellets. pH and nitrate levels were measured weekly using a WP-91 pH meter (TPS) and a HI97728 nitrate photometer (Hanna Instruments), respectively. Fertilized eggs (at the cleavage period of development) were collected in the morning and then placed in Petri dishes filled with Danieau’s solution (17 mM NaCl, 2 mM KCl, 0.12 mM MgSO_4_, 1.8 mM, Ca(NO_3_)_2_, 1.5 mM HEPES) in DSI-060D incubators (Digisystem) at either control (28°C) or elevated (30°C) temperature. Temperature inside the incubators was adjusted every 60 seconds. We selected this control temperature as it is the optimal rearing temperature of zebrafish in laboratory settings and chose the treatment temperature as +2°C as current IPCC scenarios estimate that because of climate change temperatures will globally increase between 1.5°C and 2°C above pre-industrial levels by the end of the century (Allen et al., 2018). The embryo medium was changed daily. Embryos were reared under a 14/10h light-dark cycle and from five days post fertilization (dpf) onwards they were fed a larval diet (Zeigler Bros) daily, until seven dpf. Temperatures between treatments differed significantly (Wilcoxon rank sum test, p-value < 0.001) and did not follow a normal distribution (Shapiro-Wilk test, p-value < 0.001): the “treatment” temperature over the experimentation period was 30.1 ± 0.3 °C, whereas the mean control temperature was 28.2 ± 0.3 °C (SFig. 1). This study was carried out in approval of the Committee on the Use of Live Animals in Teaching and Research (CULATR) of the University of Hong Kong (# 5504-20).

### Behavioural assays and analyses

To test the innate response of zebrafish larvae to olfactory cues, we conducted a first experiment referred to as “the innate experiment”. Larvae were exposed to Conspecifics Alarm Cue (CAC), a substance that is known to trigger an alarm response such as reduced locomotion or increased freezing, which consists in a total suppression of the swimming activity (Jesuthasan & Mathuru, 2008; Speedie & Gerlai, 2008). The catfish cue (Catfish C) was chosen to assess the larval response to an external odour from a non-predatory species, not signalling any danger in control thermal conditions (Lucon-Xiccato et al., 2020). The CAC was produced by sacrificing zebrafish larvae reared together with the experiment subjects by head concussion, and homogenizing their bodies in water at a concentration of one donor larva/mL with a sterile mortar and pestle, 5 min before cue exposure time (Lucon-Xiccato et al., 2020). For Catfish C, water in which two shark catfish were maintained for 24h (*Pangasianodon hypophthalmus*; length:water ratio of 20 cm:7L) was used following the methodology of the same study. Seven dpf larvae were transferred to experimental acrylic chambers (40 × 40 mm each) with a transparent bottom filled with six mL of Danieau’s solution. Larvae were given 40 min acclimation time, then recorded from above using a Canon EOS M50 camera. After a baseline observation period of 12 min, larvae were exposed to either CAC (n = 26 for control temperature, n = 27 for treatment temperature), Catfish C (n = 29 for control temperature, n = 28 for treatment temperature) or control water (n = 30 for control temperature, n = 27 for treatment temperature) for another 12 min (Fig. 1a). Each cue was introduced in the chamber at a volume of 0.5 mL, using a 2.5 mL syringe.

**Figure 1:**
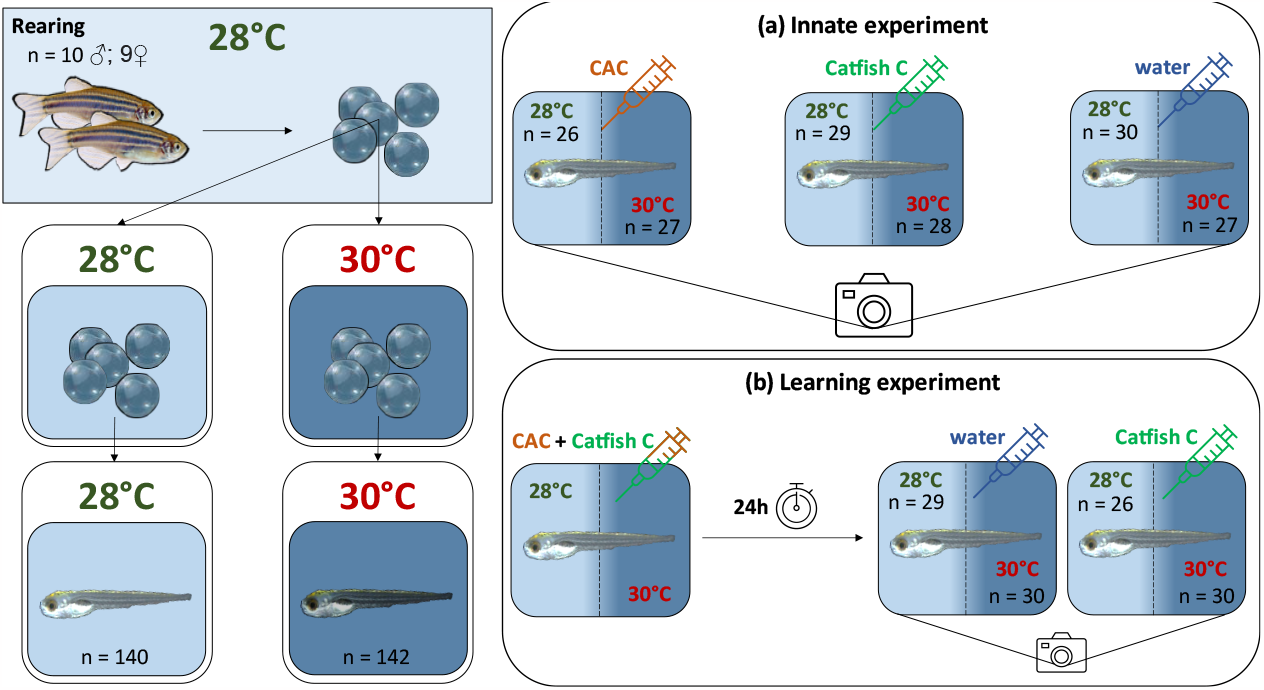
**Experimental designs** of the (a) innate and (b) learning experiments; “n” values correspond to the number of individuals within each group; treatments are illustrated by a syringe which colour corresponds to the specific cue the larvae are exposed to (Conspecifics Alarm Cue, CAC, or catfish cue, Catfish C, or water control); in the learning experiment, conditioning consists in exposing the larvae to a mixture of CAC and Catfish C.

To test whether seven dpf zebrafish can learn to associate the non-alarming catfish cue with a dangerous signal and whether temperature has an influence on that response, we conducted a second experiment referred to as “the learning experiment”. Larvae were introduced at six dpf to the experimental acrylic chambers, acclimated for 40 min and then conditioned to 1 mL of a mixture of CAC (0.5 mL) and Catfish C (0.5 mL), prepared as described above. The conditioning phase lasted 40 min, then the larvae were placed back into the incubation chambers for 24 hours, following the methodology of a previous study (Lucon-Xiccato et al., 2020). On the following day, larvae were placed back in individual chambers, where they were given 40 min to acclimate and were then video recorded. After a baseline activity period of 12 min, larvae were exposed to either control water (n = 29 at control temperature; n = 30 at elevated temperature) or Catfish C alone (n = 26 at control temperature; n = 30 at elevated temperature) for another 12min, as in experiment 1 (Fig. 1b). Larvae from both experiments were immediately snap frozen in liquid nitrogen after the behavioural assays and stored at -80°C for further processing.

To standardize the length of all videos to 12 min, raw videos were cut in Adobe Premiere 2019. The videos were then processed in DeepLabCut v 2.2.0 (Mathis et al., 2018) for body coordinates tracking of multiple animals per video. For a training dataset for machine learning of a network (Max. it. = 50 000; Saved it. = 5000; kept snapshots = 10) body parts were manually labelled in 20 frames per video among a representative set of 10 videos across dates and treatment. Incorrect tracklets were manually corrected post-analysis in DeepLabCut and positions through time of each larva were extracted in R v 4.2.1 (R Core Team, 2018) using the trajr package v 1.4.0 (McLean & Skowron Volponi, 2018). Trajectories in which position was retrieved in less than 80% of all frames were discarded (Table S1). With the filtered data, whole distances swam for each larvae every minute in each video were calculated (Table S2). Finally, this value was statistically compared in R for each larva: the values before exposure to the cues were paired to those of after exposure. The paired values were statistically compared using a paired t-test when the distribution was normal, or else with a Wilcoxon signed-rank test, to assess whether the cue influenced the behaviour. Finally, the length of each larva was measured using the software JMicroVision version (Roduit, 2004): a random frame of each raw video per experiment was extracted and the one-dimensional scale was set to 40 mm of the behavioural chamber. This allowed checking other factors such as body length potentially driving differences in swimming performance.

### RNA sequencing and gene expression analyses

In order to assess the molecular response of zebrafish to olfactory cues and the potential influence of temperature, total RNA was extracted from whole larvae using a the TRIzol™ Plus RNA Purification Kit with Phasemaker tubes (Thermo Fisher) due to the small size of the larvae. RNA concentration was measured using Qubit™ RNA HS Assay Kit (Thermo Fisher Scientific) and quality assessed using TapeStation (Agilent). RNA was sequenced at 150 bp paired end on an Illumina NovaSeq at the Centre for PanorOmic Sciences (CPOS) of the University of Hong Kong. After sequencing, raw sequence data (on average 32,022,942 ± 3,345,742; Table S3) were trimmed for adapters and filtered based on read quality using Trimmomatic (Bolger et al., 2014) with the following parameters: “ILLUMINACLIP: all_adapters.fa:2:30:10: :TRUE LEADING: TRAILING:3 SLIDINGWINDOW: : 0 MINLEN:3 “. High quality reads (on average 29,879,111 ± 3,122,986; Table S3) were then mapped against the reference genome (Genome Reference Consortium z11) from Ensembl (K. Howe, 2020), but using the more comprehensive zebrafish transcriptome annotation from Lawson et al., 2020. Mapping was done using the program HISAT2 (Kim et al., 2019) to obtain expression levels across the genome and 15,691 genes out of 36,351 mapped genes were associated to GO terms in the Lawson annotation of the GRCz11 zebrafish genome (Table S4). For functional enrichment analysis, the functional annotation file retrieved from the Lawson lab website that associated the gene IDs with ZFIN IDs (Howe et al., 2013), Ensembl IDs and/or Entrez IDs (Lawson et al., 2020), and from a custom reference set file created using BioMart resources (containing Ensembl IDs, ZFIN IDs and Entrez IDs) and the R package “FindMyFriends” (Pedersen, 2016).

Finally, differential expression analyses were led using DESeq2 v 1.38 (Love et al., 2014) to investigate which genes are differentially expressed (DE) between control larvae and treatment larvae between samples from the same experiment (innate or learning). The larvae’s date of fertilization (df) was found to be an important factor influencing the differential expression of genes, so it was kept in the design formula of both experiments (Likelihood Ratio Test, “df” accounted for respectively 1 .3% and 51.8% of the DE genes in experiment 1 and 2). The formula contained a combination of the two factors (temperature and cue) with therefore six possible levels, allowing to make specific pairwise comparisons between the most specific groups of the experimental design (“ ∽ df + combination”). Differentially expressed genes with a baseMean under 10 and/or an absolute value of log2foldchange inferior to 0.3 were discarded to ensure that differential expression was not an artifact of low counts, and to increase stringency.

For the significantly differentially expressed (DE) genes, functional enrichment analyses were performed in OmicsBox v 1.4.11 (Fisher’s Exact Test). The GO terms get associated with the gene IDs because of their correspondence with either one of the three other types of IDs: if there was no documented Ensembl ID, the Entrez ID would be used and if the two former ones were absent, then the ZFIN ID would be used. The Gene Ontology (GO) terms with an FDR adjusted p-value (Benjamini & Hochberg, 1995) below the 0.05 threshold were considered enriched and reduced to most specific.

## Results

### Behavioural response to olfactory cues

We hypothesized that warming would alter the swimming behaviour of zebrafish larvae. There was a significant effect of rearing temperature on baseline swimming distance in the innate experiment (Wilcoxon test, p-value < 0.001), with larvae swimming on average 1,127.2 ± 663.2 mm if reared in control temperature (n = 67) but only 584.7 ± 262.5 mm when reared at elevated temperature (n = 62), during the baseline activity period (Table S2 & Fig. 2). Although one individual reared at control temperature swam more distance than the others from the same group, the observed difference was still significant in its absence. Similarly, rearing temperature had a significant effect on baseline swimming distance in the learning experiment involving conditioned larvae (Wilcoxon test, p-value < 0.005), with larvae swimming on average 1,241.9 ± 771.8 mm when reared at control temperature (n = 50), but only 853.7 ± 600.1 mm if reared at elevated temperature (n = 47; Table S2 & Fig. 2). This provides two independent measures in separate experiments revealing an effect of elevated temperature on the routine swimming distance of zebrafish larvae.

**Figure 2:**
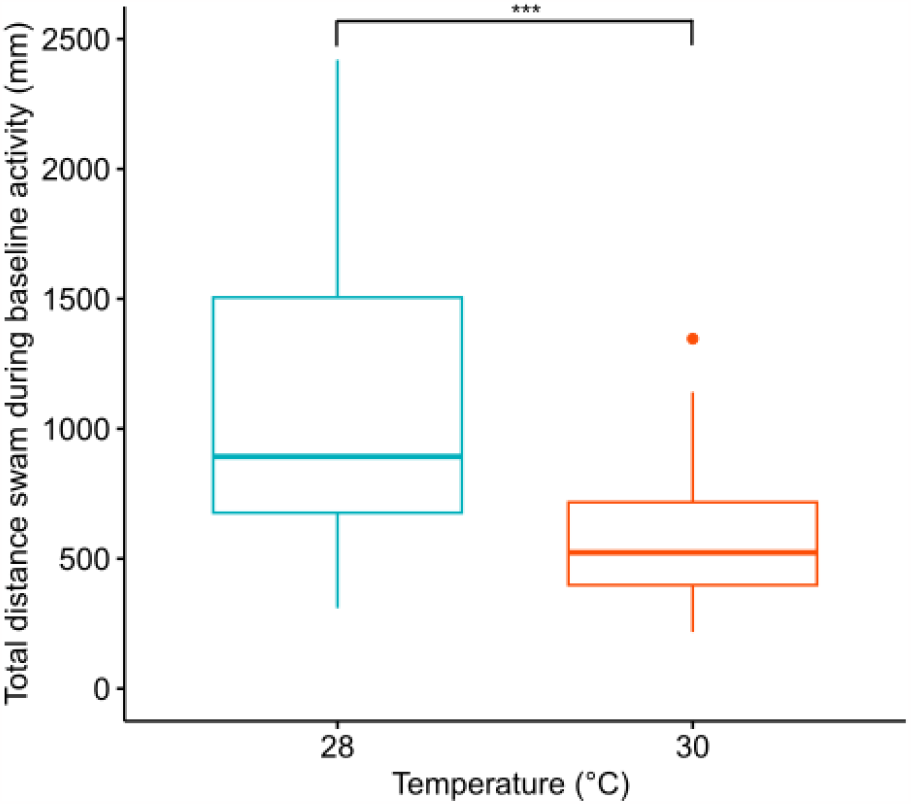
**Effect of elevated temperature on routine swimming activity** as total distance swam by the larvae of the innate experiment during their baseline activity period in control (28°C; green) or treatment (30°C; red); dots represent outliers and one outlier of the control group was removed (distance = 4000 mm); stars (***) indicate the significant difference between mean values.

Overall, there was no significant innate effect of any olfactory cue exposure on swimming activity, no matter the temperature at which they were reared (Tables S1 & S2, Fig. 3). Although not significant, there was a slight increase in the average distance swam after exposure to any cue in the innate experiment (Fig. 3a), due to individuals showing variable swimming activity (Fig. 3b, c and d). In control thermal conditions, control water did not affect the total distance swum by naive larvae, as expected, with larvae swimming 1,221.8 ± 815.6 mm during baseline period and 1,258.2 ± 790.8 mm once exposed to control water (n = 24, Wilcoxon test, p-value = 0.782; Fig. 3b). Exposure to catfish cue did not reduce the total distance swum either (n = 24, Wilcoxon test, p-value = 0.291; baseline distance = 1,013.6 ± 565.9 mm, exposure distance = 1,206 ± 659.4 mm; Fig. 3c), nor did exposure to CAC (n = 19, paired t-test, p-value = 0.545; baseline distance = 1,151 ± 569 mm, exposure distance = 1,203.8 ± 620.5 mm; Fig. 3d). Similar results were found at elevated temperature: control water did not affect the total distance swam by naive larvae (n = 22, Wilcoxon test, p-value = 0.526; SFig. 2). Larvae exposed to control water swam on average 602.7 ± 294.6 mm during the baseline period and 643.5 ± 337.2 mm during the exposure period. Exposure to catfish cue did not reduce the total distance swum either (n = 20, Wilcoxon test, p-value = 0.183; baseline distance = 559.7 ± 262.1 mm, exposure distance = 727.4 ± 427.1 mm). Finally, exposure to CAC did not have a significant effect on the total distance swum (n = 19, Wilcoxon test, p-value = 0.474; baseline distance = 590.5 ± 234.3 mm, exposure distance = 679.1 ± 333 mm).

**Figure 3:**
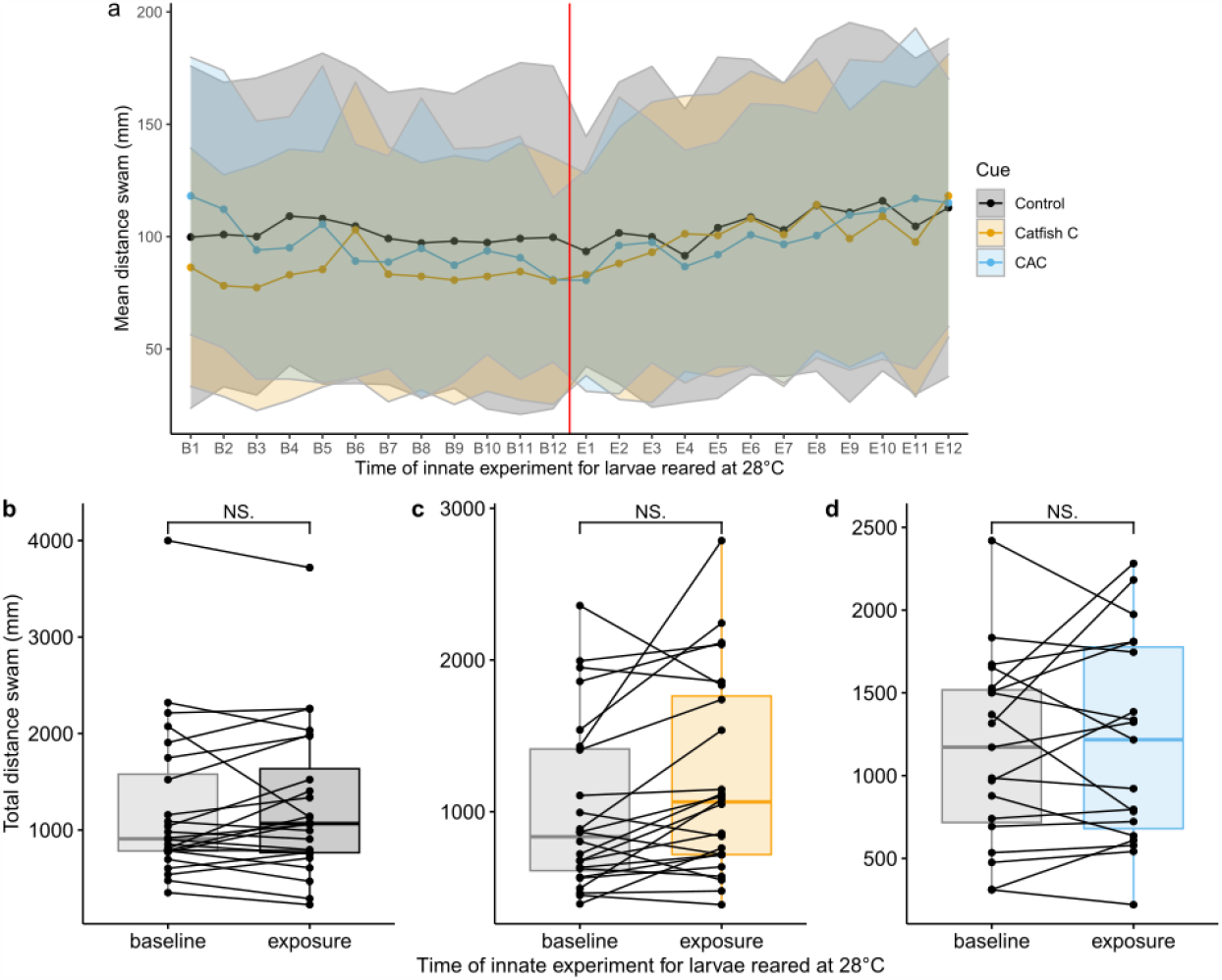
**Effect of elevated temperature on swimming response to olfactory cue exposure** as mean distance per minute (a) and total distance (b, c and d) swam by larvae reared in control temperature, before cue exposure (B; baseline) and after cue exposure (E; exposure) for each cue group: control water (C, black), catfish cue (Catfish C, orange) or Conspecifics Alarm Cue (CAC, blue); ribbons around solid lines correspond to standard deviation; black dots linked by full lines represent paired individual comparisons of before and after the cue exposure; “NS.” stands for “non-significant” and illustrate the absence of a statistical difference between the mean values.

Classical conditioning and subsequent cue exposure did not have a significant effect on the swimming activity of conditioned larvae in the learning experiment (Table S2, SFig. 3 & 4). At control temperature, exposure to control water did not significantly change the total distance swum by conditioned larvae (n = 29, Wilcoxon test, p-value = 0.422; baseline distance = 1,300.9 ± 734.8 mm, exposure distance = 1,136.1 ± 676.8 mm). No reduction in swimming distance was found with catfish cue exposure despite the previous conditioning with mixed catfish cue and CAC (n = 21, Wilcoxon test, p-value = 0.584). Larvae swam 1,262.7 ± 757.4 mm during the exposure period, which is on average 8.8% more than before smelling the cue (baseline distance = 1,160.4 ± 831.5 mm). Similar results were found at elevated temperature, as distance swum by conditioned larvae exposed to control water did not change significantly (n = 24, Wilcoxon test, p-value = 0.814; baseline distance = 831.3 ± 531.2 mm, exposure distance = 864 ± 535.2 mm) neither did the distance swum by conditioned larvae exposed to catfish cue (n = 23, Wilcoxon test, p-value = 0.155; baseline distance = 877.2 ± 675.9 mm, exposure distance = 1,094.4 ± 726.7 mm). We made sure that a number of other factors did not significantly influence routine locomotory behaviour during the baseline activity period of both experiments, such as body length (3.4 ± 0.3 mm, Wilcoxon rank sum test, p-value = 0.067), experimental chamber position (Kruskal-Wallis test, p-value = 0.725), holding aquarium tank (Kruskal-Wallis test, p-value = 0.912), or the cue administered during baseline period (Kruskal-Wallis test, p-value = 0.478). Finally, we also made sure well position and type of cue were not covariant (Pearson’s χ^2^ test, p-value = 0.570).

### Molecular responses to olfactory cues

Being reared at elevated temperature resulted in eight genes being differentially expressed compared to control temperature, in larvae not exposed to any olfactory cues. All those genes were upregulated (Table S5). Among them, three genes, *GH3 domain containing, mitochondrial ribosomal protein S9* and *threonyl-tRNA synthetase 2, mitochondrial* (*ghdc, mrps9* and *tars2*), are involved in peptide synthesis, while the gene *protocadherin 2 alpha b2* (*pcdh2ab2*) codes for a subunit of protocadherin protein, that is involved in neuronal development.

At control temperature, exposure to cues resulted in no gene being differentially expressed in whole zebrafish larvae exposed to CAC, and only in one upregulated gene, *pcdh2ab2*, in larvae exposed to catfish cue (Table S6). At elevated temperature, the *mitochondrial ribosomal protein S6* (*mrps6*; Table S7) only was downregulated in larvae exposed to CAC. However, a much larger response with 740 differentially expressed genes (Table S6) was found for catfish cue at elevated temperature, with a variety of altered key functions, such as ATP metabolism processes, protein synthesis processes, cell signalling and neurotransmission.

Among the enriched processes in larvae that developed at elevated temperature when exposed to catfish cue, cellular organisation and localization were upregulated (Table S8). Specifically, a large majority of genes involved in GTPase activity were over-expressed, enriching small GTPase mediated signal transduction and GTPase regulator activity (Table S8). Proteins encoded by these genes were either involved in the activation of GTPases or were GTPases themselves involved in signal transduction, such as Rho small GTPases or Rab protein (*rab8a)* and other types of G-protein, such as the alpha activating activity polypeptide, olfactory type (*gnal*). Furthermore, palmitoylation of proteins was also upregulated, enriching lipoprotein metabolism (Table S8), due to upregulation of genes such as *zdhhc12b* and *zdhhc5b*, coding for zinc finger DHHC-type palmitoyltransferases, and *golga7* and *golga7ba*, coding for golgins 7A proteins part of the palmitoyltransferase complex (Table S9).

Finally, genes involved in neuronal development processes were upregulated (Table S8) when larvae responded to catfish cue at elevated temperature, notably through the differential expression of genes involved in axon guidance, in particular of olfactory sensory neurons (*ntn1a* and *unc5b*), contactin-associated genes (*cntnap2a* and *cntnap5a*) involved in organization of myelinated axons as well as axon growth associated genes of the semaphorin-plexin pathway (*sema4ab, plxna1a* and *plxna2)*. Additionally, genes involved in synaptic transmission were over-expressed upon exposure to catfish cue at elevated temperature. These genes belong to calcium, sodium and potassium voltage-gated ion channels (Table S9), leading to functional enrichments in metal ion binding and voltage-gated potassium channel activity (Table S8). Furthermore, γ-aminobutyric acid (GABA) receptor genes (*gabrg2, gabrg3* and *gabrz*; Fig. 4a; Table S9) were also upregulated.

**Figure 4:**
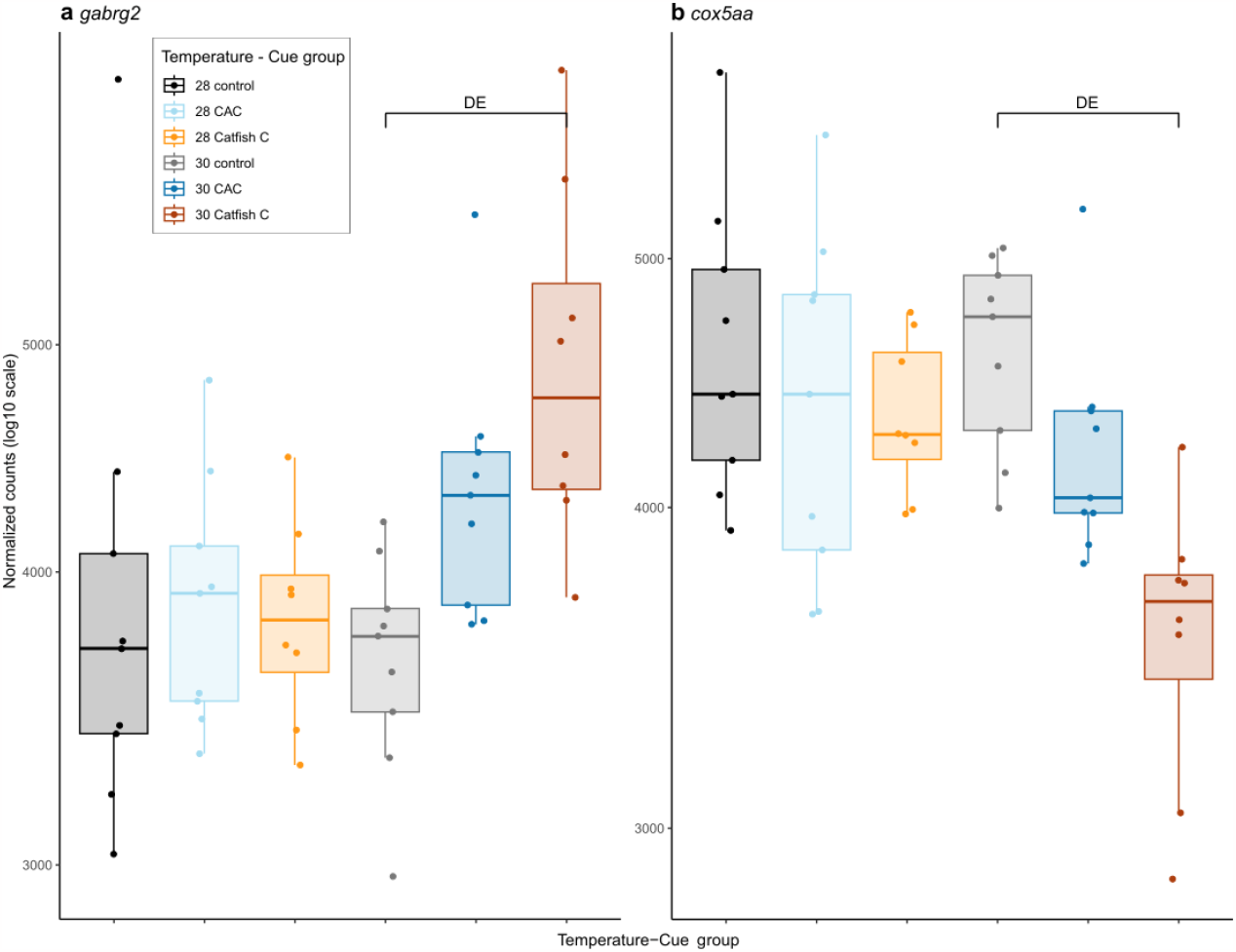
**Effects of olfactory cue exposure and elevated temperature on gene expression** as normalized counts (log_10_) in all temperature-cue groups for the gene cox5aa, a subunit of the cytochrome c oxidase (a) and for the gene gabrg2, a subunit of GABA_A_ receptor (b); larvae were grouped as a function of the temperature they were reared at (control, 28 or elevated, 30) and the cue they were exposed to (control, C; Conspecifics Alarm Cue, CAC; catfish cue, Catfish C); each dot represents an individual; “DE” shows which pairwise comparison revealed this gene as differentially expressed.

We also found many energy-related processes altered (Table S9), such as proton transmembrane transport, mitochondrial electron transport and organization and mitochondrial ATP synthesis. This is due to the downregulation of genes involved in respiratory complexes of the electron chain (Table S8) including the NADH:ubiquinone oxidoreductase, the ubiquinol-cytochrome c reductase, the cytochrome c oxidase (Fig. 4b) and the ATP synthase (Table S8). Protein translation and homeostasis also exhibited reduced transcription upon exposure to catfish cue at elevated temperature. For example, several processes within protein synthesis were altered, including seryl-tRNA aminoacylation, rRNA binding and structural constituents of ribosome (Table S8). The same was found for protein folding, a post-translational mechanism, as two genes coding subunits of the chaperonin containing TCP1 protein (*cct2* and *cct4*) were downregulated (Table S9).

In the learning experiment, no differentially expressed genes were found in conditioned zebrafish exposed to catfish cue regardless of temperature.

## Discussion

In this study we explored how temperature, specifically predicted near-future warming, affects the innate olfactory response in zebrafish larvae and the underlying molecular drivers. We found that exposure to elevated temperature during embryogenesis until seven days post fertilization (dpf) decreased routine swimming distance and increased expression of genes involved in peptide synthesis and neuron development. However, unexpectedly, none of the cues provoked a reduction in swimming activity, no matter the rearing temperature, despite elevated temperature mediating a large molecular response upon exposure to non-alarming catfish cue. Finally, we found that conditioned larvae did not react to catfish cue in a way consistent with associative learning neither behaviourally nor molecularly.

Elevated temperature during embryogenesis resulted in a reduction in swimming distance in zebrafish larvae. As previous findings found no swimming alteration when seven dpf zebrafish reared at control temperature were suddenly exposed to 30°C (Abozaid et al., 2020), our results suggest that reduced swimming activity is an effect of developmental thermal exposure. Reduced swimming could be a behavioural trade-off caused by higher energy costs of maintaining homeostasis at 30°C, as the thermal performance curve dogma predicts (Huey & Stevenson, 1979). Concurrently, elevated temperature triggered an over-expression of genes involved in neuron development and protein synthesis. In particular, *pcdh2ab2*, which codes for a neuronal cell-surface protocadherin expressed throughout the developing nervous system (Emond & Jontes, 2008; Tada et al., 2004), is upregulated in larvae at elevated temperature. Similarly, genes such as an acid-amino acid ligase (*gdhc)*, a ribosome constituent *(mrps9)* and *tars2*, which codes for an enzyme that performs the first step of translation, are found at elevated levels in elevated temperature. Upregulation of genes involved in peptide production and brain development at elevated temperature is consistent with the acceleration of metabolism and overall development due to high temperature (López-Olmeda & Sánchez-Vázquez, 2011). Moreover, a 2°C elevation was also reported to increase larval expression of growth hormone and insulin-like growth factor genes in other fish species (Politis et al., 2017). Therefore, our results support that larvae exposed to 30°C during embryogenesis had a faster development regarding molecular patterns, than larvae reared at control temperature.

While elevated temperature elicited a behavioural change, neither of the cues provoked freezing behaviour, no matter the rearing temperature. A complete suppression of locomotory movement has been previously reported in response to alarming olfactory substances in zebrafish (Jesuthasan & Mathuru, 2008; Lucon-Xiccato et al., 2020; Norton & Bally-Cuif, 2010), which is likely a survival strategy to remain hidden from a predator. Therefore an absence of behavioural and molecular response to CAC is contrary to our expectations, as injured conspecific cues have been shown to reduce mobility of larvae from five to 24 dpf (Jesuthasan et al., 2021; Lucon-Xiccato et al., 2020). One of the possible explanations for the lack of freezing behaviour in our samples is that larvae could have responded with behaviours other than freezing, which however, were not measured here. Indeed, freezing is not the only possible behavioural response to alarm cues and zebrafish has also been shown to react to alarming cues by bottom swimming or escaping the area (Speedie & Gerlai, 2008). Alternatively, the concentration of the cue could have been under a threshold for a behavioural (and molecular) response to occur, despite the fact that we followed the methodology of Lucon-Xiccato et al. (2020), as 7 dpf larvae are smaller than 12 dpf ones. Indeed, previous research found that intensity of the alarm response is dose-dependent (Speedie & Gerlai, 2008).

Catfish cue was used as a “neutral non-alarming cue” since the catfish (*Pangasianodon hypophthalmus*) is not a known predator to zebrafish and its smell has not been found to trigger a freezing response (Lucon-Xiccato et al., 2020). The absence of freezing behaviour in response to catfish cue is therefore consistent with our expectations and with previous research. Interestingly, exposure to catfish cue elicited expression changes of *pcdh2ab2* at control temperature. In seahorses (*Hippocampus erectus*), *pcdh2ab2* is also changed in expression levels when faced with the visual and olfactory signal of a paired mate (Mederos et al., 2022). As *pcdh2ab2* is involved in neuron development, it could play a role in the neuronal alterations in the context of odour detecting and olfactory memory. In our case, since this gene was not also upregulated in larvae exposed to CAC, the reason for upregulation of this gene upon smelling catfish cue might be due to the novelty of this specific cue in the larvae’s environment. Unlike CAC, which was prepared with larvae of the same school, catfish cue was never introduced before and this was the first exposure to this new smell for the larvae.

Elevated temperature mediated a large transcriptional response to the novel smell of catfish cue exposure. Processes involved in cellular signalling, cell organisation and localization were upregulated, including genes involved in GTPase activity essential for signal transduction, particularly in G-proteins signalling pathways. Small G-protein coding genes of the Rho family participate in cell shaping (Csépányi-Kömi et al., 2012), while genes coding for Rab proteins like *rab8a* are involved in the elongation of sensory cilia (Omori et al., 2008), which are cell organelles harbouring olfactory receptors (Singla & Reiter, 2006). Interestingly, the olfactory specific G-protein gene *gnal* was upregulated as well, which codes for an olfactory specific G-protein expressed in the ciliated olfactory sensory neurons of zebrafish and is involved in olfactory map refinement in the olfactory epithelium (Dang et al., 2018; Yoshihara, 2014). Similarly, odorant exposure in mice leads to transcriptional changes in cell signalling and G-protein-coupled receptor activity in olfactory sensory neurons (Horgue et al., 2022) and novel odour in particular provokes expression changes in cytokine-mediated cell signalling genes in rats (Irwin & Byers, 2012; Montag-Sallaz & Buonviso, 2002). Similar processes therefore seem to occur in fish after detecting a novel odour in the environment at high temperature, showing that there is a change in signalling activity possibly involved in neuronal olfactory circuitry modifications in response to novel odours.

Together with cell signalling, genes involved in neuronal development were also upregulated in response to catfish cue at elevated temperature. In particular, *ntn1a* and *unc5b* participate in sensory axon targeting in the olfactory bulb (Dang et al., 2023; Lakhina et al., 2012). Moreover, over-expression of contactin-associated genes, such as *cntnap2a*, required for proper organization of myelinated axons (Poliak et al., 1999), suggests that the cellular organization of neurons is modified after catfish cue exposure. Finally, genes of the semaphorin-plexin pathway participate in axon growth, notably of neurons of the olfactory system (Emerson et al., 2018; Marcos et al., 2017), further supporting the hypothesis that detection of novel odours at high temperature triggers neuronal growth. Other upregulated genes participated in general neurotransmission, such as voltage-dependant ion channels and neurotransmitter receptors like GABA_A_ receptors. The reason for such a strong response to catfish cue at elevated temperature only might again lie in thermally-induced accelerated development, in particular of the nervous system, with larvae that developed at elevated temperature potentially having more neurons and/or synapses at elevated temperature compared to same age larvae reared at control conditions (López-Olmeda & Sánchez-Vázquez, 2011). Along with accelerated development, warming during the larval stage could also provoke changes to neuronal circuitry resulting in altered olfactory detection. This is the case of honeybees, for example, in which the olfactory-input region of the brain is altered due to differences in developmental temperature (Groh et al., 2004). The upregulation of axon guidance genes in particular supports both hypotheses, as it would allow more axons to project into the olfactory bulb. Whether through accelerated development, altered olfaction neural circuitry or a combination of both, temperature has therefore been found here to strongly impact novel odour recognition in fish at the molecular level, suggesting neuromolecular changes in a future warming world.

On the contrary, different genes coding for components of the electron transport chain were downregulated. Downregulation of electron transport chain genes is expected to reduce mitochondrial respiration, which could relate to hypometabolism under heat stress, as previously seen in mitochondria of other fishes at elevated temperature (Chung & Schulte, 2015; Michaelsen et al., 2021). Genes involved in protein synthesis were also downregulated, which further indicates the repression of several metabolic processes due to temperature-induced increased metabolism. A possible explanation to why this type of response to catfish cue is only observed at elevated temperature could be that detecting new odours while maintaining homeostasis at high temperature requires additional energy. This therefore creates a trade-off between expensive basal processes, such as protein production, and novel odour response. Since protein synthesis is one of the most expensive cellular processes, it is also one of the first ones that is repressed (Advani & Ivanov, 2019), here to redirect energy to the odour response Consequently, when a novel odour is detected, energetic reserves would shift towards olfactory responsive functions, possibly leading to energy resource depletion and long-term detrimental effects. Although this large metabolic repression is observed after exposure to catfish cue, at elevated temperature CAC also triggered downregulation of the ribosomal protein coding *mrps6* gene, which could indicate that energetic trade-offs are necessary to respond to several types of olfactory cue at elevated temperature. Overall, our results therefore show how future-predicted temperature exposure during development mediates increased expression of genes involved in cell communication, neuron functioning and development, at the cost however of a repression of key metabolic processes following an otherwise non-noxious environmental stimulus that is the catfish cue. Such energy reallocation needed during processing an environmental signal might negatively affect basic processes necessary to development and growth in the long run, when faced with future global warming conditions.

Conditioning did not result in any behavioural or molecular response, with conditioned larvae not exhibiting freezing behaviour or gene expression changes following catfish cue exposure regardless of temperature. This was unexpected, since older 24 days post-fertilization (dpf) larvae respond to catfish cue by freezing after conditioning with CAC (Lucon-Xiccato et al., 2020), and some larvae of similar age to ours also demonstrate associative learning through classical conditioning, although to visual stimuli and not olfactory cues (Pritchett & Brennan, 2020). Despite these contradictory results, the possibility that seven dpf larvae are not developed enough for robust learning cannot be discarded, as the target performance in previous associate learning studies were modest and not always consistent across all larvae (Gerlai, 2016; Pritchett & Brennan, 2020). Regarding the absence of gene expression changes upon smelling catfish cue in conditioned larvae, these findings reinforce our hypothesis of the novel odour stress response at high temperature. Contrary to the innate experiment, where the large molecular response to the catfish cue was seen when larvae had their first exposure to it, in the conditioning experiment larvae were tested for a response to the catfish cue based on their previous experience with no large molecular reprogramming. In summary, our findings show an effect of future-predicted thermal conditions on developing zebrafish, causing reduced swimming during routine activity and largely altering gene expression processes in the nervous system as larvae encounter a new olfactory stimulus, making environmental sensing energetically costly.

## Data availability

The raw sequencing data can be found in BioProject PRJNA974200. The reviewer link to the data is: https://dataview.ncbi.nlm.nih.gov/object/PRJNA974200?reviewer=n5uqu49k1qf7jj3ljqijee7oj8

## Author contributions

JMS conceived and carried out the experiments with input from CS. JS provided the breeding animals and input on zebrafish breeding. JMS analysed the data under the supervision of CS and LCB. JMS wrote the paper with input from CS and LCB, and JS contributed to the final version.

## Acknowledgments

JMS and this study were funded by the start-up of CS from the University of Hong Kong. We thank Yan Chit Kam who contributed to the daily care of breeding animals and all the members of the lab for support.

