## Supplementary Data - Figures for "Warming affects routine swimming activity and novel odour molecular response in larval zebrafish"

### Supplementary Figures:

| **Name** | **Title** |
| --- | --- |
| Supplementary Figure 1 | Measured temperature (°C) regimes in each experimental group (control or elevated) throughout the experiments; stars (***) indicate the significant difference between the mean values |
| Supplementary Figure 2 | Total distance (mm) swam by the larvae reared in elevated temperature, before cue exposure (baseline) and after cue exposure (exposure) for each cue group: control water (a), catfish cue (b) or CAC (c); black dots linked by full lines represent paired individual comparisons of before and after the cue exposure; “NS.” stands for “non-significant” and illustrate the absence of a statistical difference between the mean values |
| Supplementary Figure 3 | Total distance (mm) swam by the conditioned larvae reared in control temperature, before (baseline) and after cue exposure (exposure) to control water (a) or catfish cue (b); black dots linked by full lines represent paired individual comparisons of before and after the cue exposure; “NS.” stands for “non-significant” and illustrate the absence of a statistical difference between the mean values |
| Supplementary Figure 4 | Total distance (mm) swam by the conditioned larvae reared in elevated temperature, before (baseline) and after cue exposure (exposure) to control water (a) or catfish cue (b); black dots linked by full lines represent paired individual comparisons of before and after the cue exposure; “NS.” stands for “non-significant” and illustrate the absence of a statistical difference between the mean values |

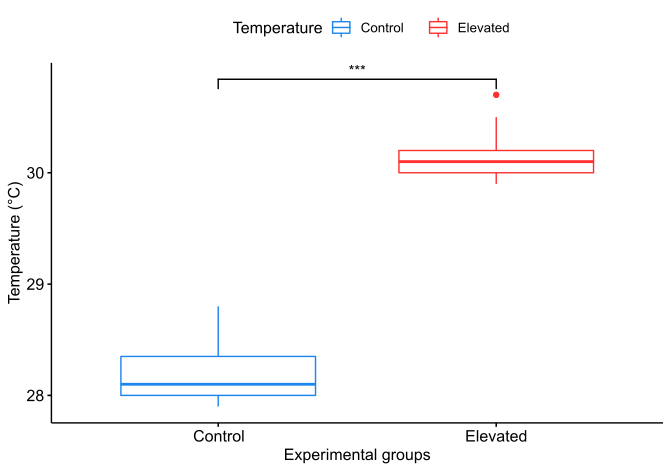
Supplementary Figure 1:

Supplementary Figure 1: Measured **temperature (°C) regimes** in each experimental group (control or elevated) throughout the experiments; stars (***) indicate the significant difference between the mean values

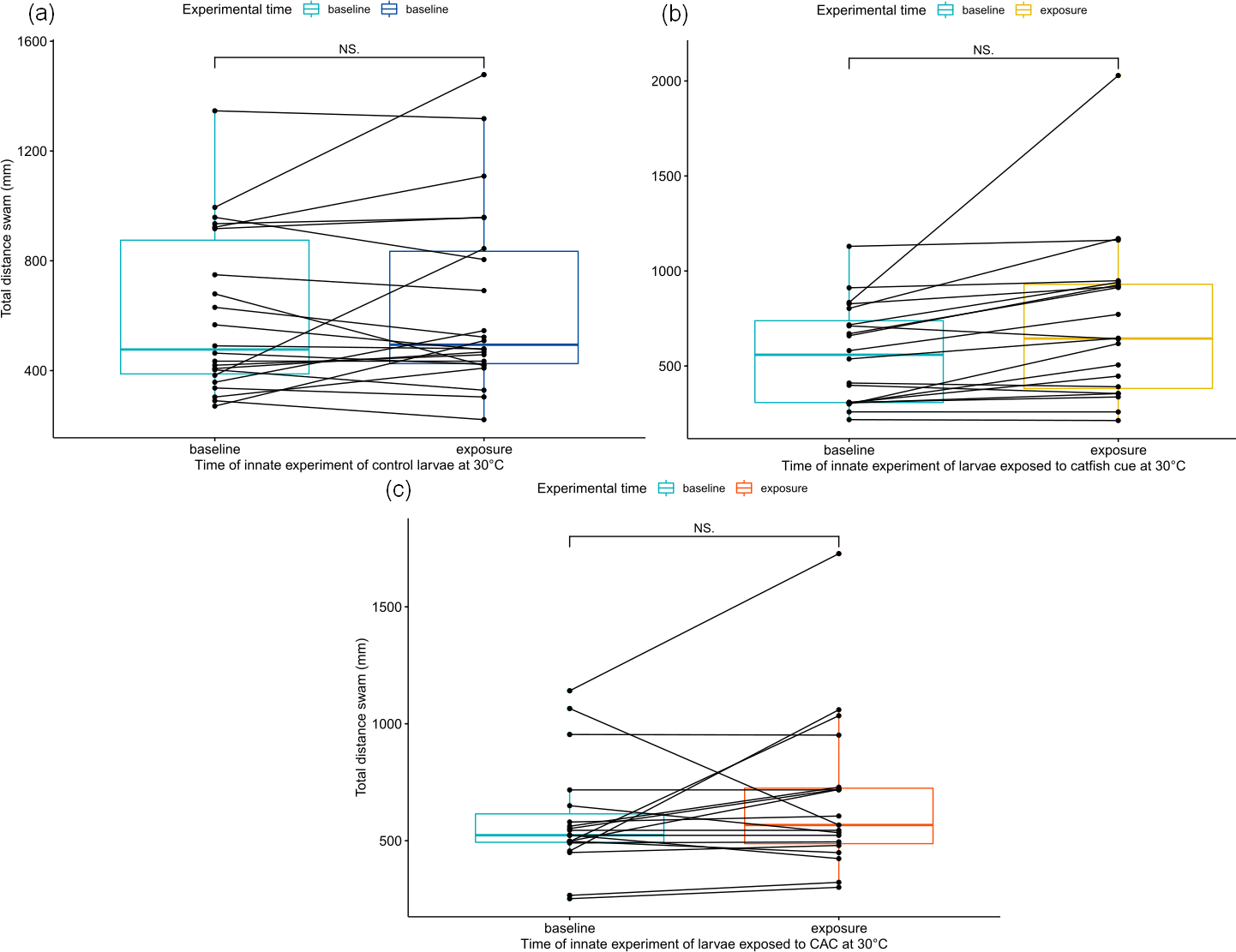
Supplementary Figure 2:

Supplementary Figure 2: Total distance (mm) swam by the larvae reared in elevated temperature, before cue exposure (baseline) and after cue exposure (exposure) for each cue group: control water (a), catfish cue (b) or CAC (c); black dots linked by full lines represent paired individual comparisons of before and after the cue exposure; “NS.” stands for “non-significant” and illustrate the absence of a statistical difference between the mean values

Supplementary Figure 3:

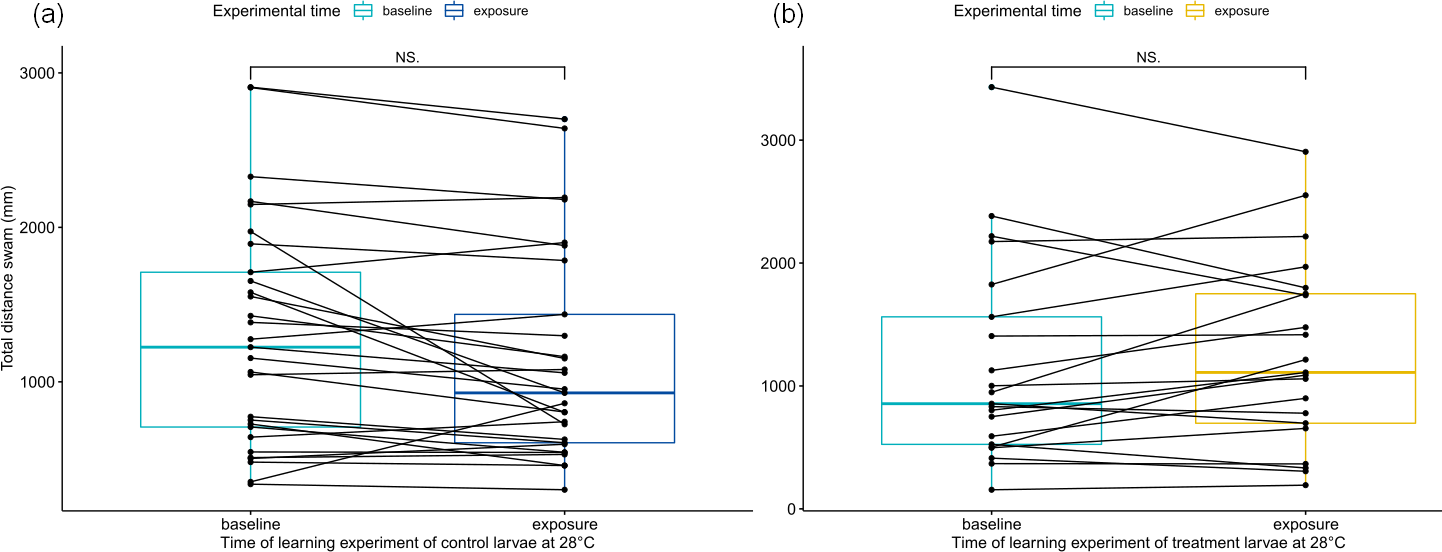

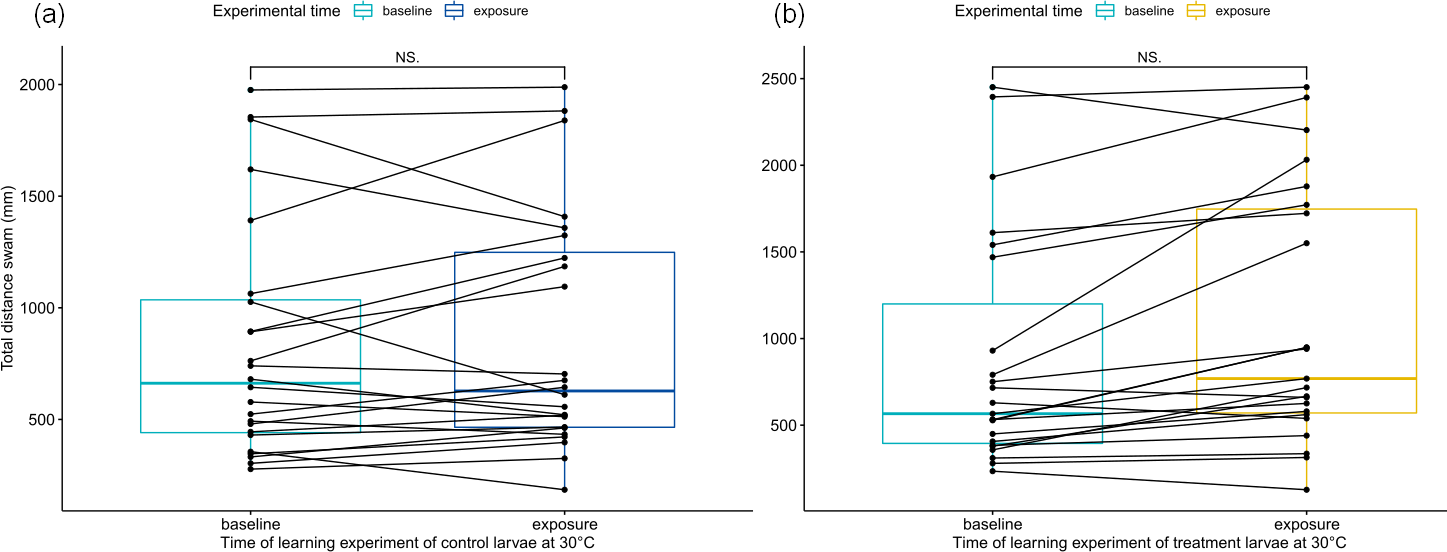
Supplementary Figure 4:

Supplementary Figure 3: Total distance (mm) swam by the conditioned larvae reared in control temperature, before (baseline) and after cue exposure (exposure) to control water (a) or catfish cue (b); black dots linked by full lines represent paired individual comparisons of before and after the cue exposure; “NS.” stands for “non-significant” and illustrate the absence of a statistical difference between the mean values

Supplementary Figure 4: Total distance (mm) swam by the conditioned larvae reared in elevated temperature, before (baseline) and after cue exposure (exposure) to control water (a) or catfish cue (b); black dots linked by full lines represent paired individual comparisons of before and after the cue exposure; “NS.” stands for “non-significant” and illustrate the absence of a statistical difference between the mean values
